# ENIGMA HALFpipe: Interactive, reproducible, and efficient analysis for resting-state and task-based fMRI data

**DOI:** 10.1101/2021.05.07.442790

**Authors:** Lea Waller, Susanne Erk, Elena Pozzi, Yara J. Toenders, Courtney C. Haswell, Marc Büttner, Paul M. Thompson, Lianne Schmaal, Rajendra A. Morey, Henrik Walter, Ilya M. Veer

## Abstract

The reproducibility crisis in neuroimaging has led to an increased demand for standardized data processing workflows. Within the ENIGMA consortium, we developed `HALFpipe` (<u>H</u>armonized <u>A</u>na<u>L</u>ysis of <u>F</u>unctional MRI p<u>i</u>p<u>e</u>line), an open-source, containerized, user-friendly tool that facilitates reproducible analysis of task-based and resting-state fMRI data through uniform application of preprocessing, quality assessment, single-subject feature extraction, and group-level statistics. It provides state-of-the-art preprocessing using `fMRIPrep` without the requirement for input data in Brain Imaging Data Structure (BIDS) format. `HALFpipe` extends the functionality of `fMRIPrep` with additional preprocessing steps, which include spatial smoothing, grand mean scaling, temporal filtering, and confound regression. `HALFpipe` generates an interactive quality assessment (QA) webpage to assess the quality of key preprocessing outputs and raw data in general. `HALFpipe` features myriad post-processing functions at the individual subject level, including calculation of task-based activation, seed-based connectivity, network-template (or dual) regression, atlas-based functional connectivity matrices, regional homogeneity (ReHo), and fractional amplitude of low frequency fluctuations (fALFF), offering support to evaluate a combinatorial number of features or preprocessing settings in one run. Finally, flexible factorial models can be defined for mixed-effects regression analysis at the group level, including multiple comparison correction. Here, we introduce the theoretical framework in which `HALFpipe` was developed, and present an overview of the main functions of the pipeline. `HALFpipe` offers the scientific community a major advance toward addressing the reproducibility crisis in neuroimaging, providing a workflow that encompasses preprocessing, post-processing, and QA of fMRI data, while broadening core principles of data analysis for producing reproducible results. Instructions and code can be found at https://github.com/HALFpipe/HALFpipe.

## Introduction

The application of neuroimaging, in particular functional MRI (fMRI), has led to an explosion in knowledge about brain functions implicated in a range of human behaviors, cognitive processes, and emotions. Such research has been spurred by rapid advances in computationally intensive software required to perform complex algorithmic processing and statistical modeling of fMRI data. The resulting proliferation of software tools designed to fulfill various analytic functions has produced a large array of options for carrying out any given type of processing. Since the fMRI signal indirectly captures the neural processes of interest, a series of computational operations on fMRI data, referred to as the *analysis pipeline*, are necessary to arrive at interpretable results. In practice, each step is flexible and subject to a number of choices by the researcher, termed *analytic flexibility* (Poldrack et al., 2017). The steps in the analysis pipeline may be reordered, run with different parameters, or may be completely omitted in some cases. Understandably, users expect different tools performing the same function to generate (near) identical results when supplied with given input data. However, the multiplicity of tools has had the unintended consequence of generating inconsistent results from studies designed to answer the same research question, sometimes even when the same data is used as the starting point (Botvinik-Nezer et al., 2020). Thus, *analytic flexibility* combined with the number of analysis steps, as well as the possible parameters for running each analysis step, has led to a vast multiplicity of methodologic variants and an equal number of possible results. This situation has contributed in part to what is widely hailed as a *crisis of reproducibility*, which now plagues the neuroimaging field (Gorgolewski et al., 2016; Poldrack et al., 2017).

One solution to improving reproducibility is to constrain the parameter space by limiting choices to default parameters established from empirically-derived best-practices (Grüning et al., 2018). For instance, established pipelines such as `fMRIPrep` (Esteban, Markiewicz, Blair, et al., 2019) and `C-PAC` (Craddock et al., 2013) have automated many of these choices. An alternative approach is to run multiple analyses separately on the same input data with the same or different pipelines, but with different parameter selections for each analysis, and then compare the results. This second approach, termed *multiverse analysis* (Steegen et al., 2016), has the advantage that results from multiple analyses may be compared and alternate solutions may be presented in published reports to promote increased transparency. However, *multiverse analysis* has the disadvantage that it may ultimately not be possible to determine the optimal or even the correct solution, as true effects in non-simulated fMRI data are often unknown.

The reproducibility crisis has led to an increased demand for standardized workflows to conduct both the preprocessing and postprocessing stages of fMRI analysis. The recent introduction and widespread adoption of standardized pipelines for fMRI data preprocessing has provided the research community with much-needed high-quality tools that have improved reproducibility (Thompson, Ching, et al., 2020). The four ingredients that are essential to data analysis and reproducible results are: (1) data and metadata availability, (2) code usage and transparency, (3) software installability, and (4) re-creation of the runtime environment. Relative to other processing pipelines, `fMRIPrep` (Esteban, Markiewicz, Blair, et al., 2019) has grown in popularity due to its adoption of best practices, open-source availability, favorable user experience, and *glass-box* principles of transparency (Poldrack et al., 2019). However, `fMRIPrep` is limited to the so-called preprocessing steps of fMRI data analysis, whilst variability in parameter selection for subsequent post-processing analysis steps (e.g., data cleaning, feature extraction, model specification) may compromise reproducibility.

The ENIGMA consortium has addressed the reproducibility crisis by pooling observational study data from structural and diffusion imaging (and more recently EEG and MEG), and by developing standardized pipelines, data harmonization methodology, and quality control protocols (Thompson, Jahanshad, et al., 2020). These workflows have successfully analyzed structural and diffusion MRI data aggregated from large numbers of small- and medium-sized cohorts to accumulate sufficient power to yield robust results on a wide range of neuropsychiatric conditions (e.g. Hoogman et al., 2020; Schmaal et al., 2020; van den Heuvel et al., 2020). However, until now the ENIGMA consortium has lacked the ability to reliably conduct consortium-wide analyses on fMRI data. More recently, however, the ENIGMA task-based (Veer et al., 2019) and resting-state fMRI (Adhikari et al., 2019) working groups have spurred initiatives to bring the ENIGMA framework to the functional domain.

To support these initiatives within ENIGMA, we developed a standardized workflow that encompasses the essential elements of task-based and resting-state fMRI analyses from raw data to group-level statistics, builds on the progress and contributions of `fMRIPrep` developers, and extends its functionality beyond preprocessing steps to include additional preprocessing, postprocessing, and interactive tools for quality assessment. These extended features include: automatic and reliable conversion of fMRI data to BIDS format, spatial smoothing, temporal filtering, extended confounds regression, calculation of task-based activations, and restingstate feature extraction, including seed-based functional connectivity, network-template (dual) regression, atlas-based functional connectivity matrices, regional homogeneity (ReHo) analysis, and fractional amplitude of low frequency fluctuations (fALFF). Although each of these post-processing functions is available in other software packages and a few pipelines have incorporated a subset of these features, `HALFpipe` combines all these post-processing tools from open-source neuroimaging packages with the preprocessing steps performed by `fMRIPrep` (see Table 1). Furthermore, although `HALFpipe` provides recommended settings for each of the processing steps (see Table 2), it allows users to run any combinatorial number of these processing settings, thereby offering a streamlined infrastructure for pursuing multiverse analyses. Similar to other processing pipelines, `HALFpipe` is available as a containerized image, thereby offering full control over the computational environment. In this article, we provide a detailed description of `HALFpipe`. First we explain the software architecture and implementation, followed by a walkthrough of the procedure for running the software, and finally a discussion of the potential applications of the pipeline.

**Table 1:** Comparison to other neuroimaging pipelines

|  |  | HALFpipe | C-PAC | fMRIPrep<br>MRIQC<br>FitLins | CONN<br>toolbox | XCP<br>Engine | DPARSF<br>DPABI |
| --- | --- | --- | --- | --- | --- | --- | --- |
| Quality<br>assessment | Quality metrics | Yes | Yes | Yes | Yes | Yes | Yes |
|  | Visual quality<br>assessment | Yes | Yes | Yes | Yes | Yes | Yes |
| Features | Task-based<br>activation | Yes | No | Yes | No* | Yes | No |
|  | Seed-based<br>connectivity | Yes | Yes | No | Yes | Yes | Yes |
|  | Dual regression | Yes | Yes | No | Yes | No | Yes |
|  | Atlas-based<br>connectivity<br>matrix | Yes | Yes | No | Yes | Yes | Yes |
|  | ReHo | Yes | Yes | No | Yes** | Yes | Yes |
|  | fALFF | Yes | Yes | No | Yes | Yes | Yes |
|  | Group statistics | Yes | Yes | Yes | Yes | No | Yes |

Note: \* Task-based connectivity is supported; \*\* As implemented with LCOR (Local Correlation).

**Table 2:**
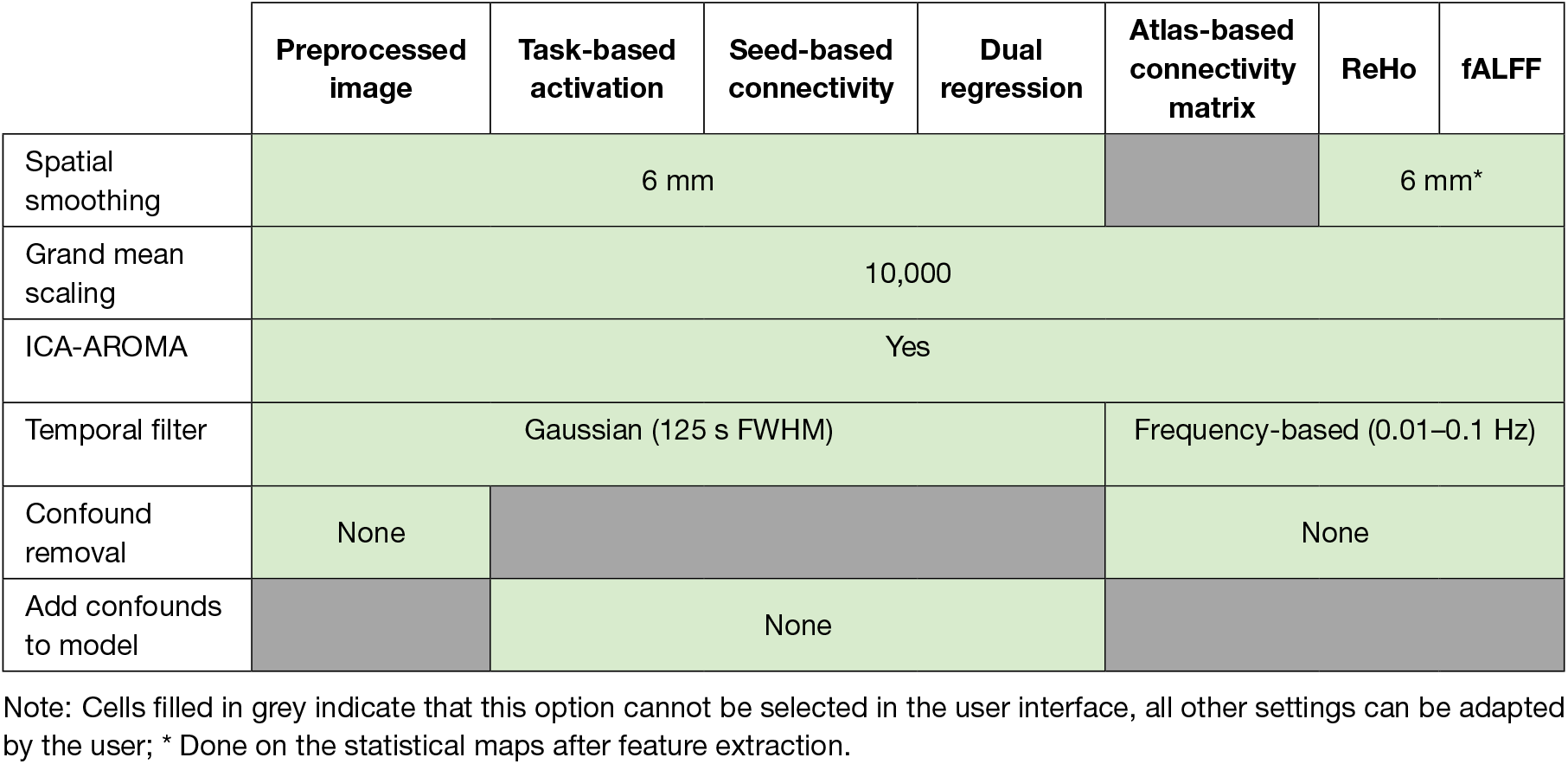
Default values for preprocessing settings per feature

|  | Preprocessed image | Task-based activation | Seed-based connectivity | Dual regression | Atlas-based connectivity matrix | ReHo | fALFF |
| --- | --- | --- | --- | --- | --- | --- | --- |
| Spatial smoothing | 6 mm |  |  |  |  | 6 mm* |  |
| Grand mean scaling | 10,000 |  |  |  |  |  |  |
| ICA-AROMA | Yes |  |  |  |  |  |  |
| Temporal filter | Gaussian (125 s FWHM) |  |  |  | Frequency-based (0.01–0.1 Hz) |  |  |
| Confound removal | None |  |  |  | None |  |  |
| Add confounds to model |  | None |  |  |  |  |  |

Note: Cells filled in grey indicate that this option cannot be selected in the user interface, all other settings can be adapted by the user; \* Done on the statistical maps after feature extraction.

## Methods

The `HALFpipe` software is containerized, similar to `fMRIPrep` or `C-PAC`. This means that it comes bundled with all other software that is needed for it to run, such as `fMRIPrep` (Esteban, Markiewicz, Blair, et al., 2019), `MRIQC` (Esteban et al., 2017), `FSL` (Jenkinson et al., 2012), `ANTs` (Avants et al., 2011), FreeSurfer (Fischl & Dale, 2000), and `AFNI` (Cox, 1996; Cox & Hyde, 1997). As such, all users of one version of `HALFpipe` will be using the exact same versions of these tools, because they come with the container. Thus, the containerization of `HALFpipe` software aids reproducibility across different researchers and computing environments. We have provided the `HALFpipe` application in a Singularity container and a Docker container. Singularity or Docker, which are both freely available, must be installed prior to downloading the containerized `HALFpipe` application. Both Docker and Singularity perform so-called operating-system-level virtualization, but are more efficient and require less resources than virtual machines. Running Docker containers on a Linux or macOS operating system requires administrator privileges. Singularity is typically run on a Linux operating system, which may be used without administrator privileges.

Besides containers, our `HALFpipe` development team adopted other software engineering best practices, which promoted faster development and reduced code errors. These industry best-practices, which have found their way into research applications (Das, 2018), involve writing code that is easy to read (albeit generally harder to write), the breakdown of complex systems into several simpler subsystems, dedicated effort toward thoughtful code design before implementation, and performing continuous integration via unit tests (Beck, 1999). The `HALFpipe` development team applied these practices whenever possible.

`HALFpipe` is being developed as an open-source project and is accepting contributions that offer new features, enhance functionality, or improve efficiency. All changes will be reviewed manually. Additionally, before inclusion in the source tree, changes will undergo automated (unit) testing, which includes running an entire analysis for one subject of the OpenNeuro dataset ds000108 (Wager et al., 2008). This way, unexpected side effects and bugs can be caught and corrected before causing problems for users.

### Databases

To automatically construct a neuroimaging processing workflow, the program needs to be able to fulfill queries such as `“retrieve the structural image for subject x”`. Many programs implement such queries using a database system.

For example, the Python implementation of the BIDS standard, called `PyBIDS`, uses an SQL database on the back-end. Each file of a BIDS dataset is assigned a number of tags in the database, such as `subject, task`, or `session` to facilitate executing queries on these tags. The PyBIDS equivalent of the example above would be `layout.get(datatype=‘anat’, sub=‘x’)`. The database approach adds a layer of complexity, because the database must be defined and populated and code must be written to interface with the database. For `HALFpipe`, we chose to construct indices and query functions based on native Python data types.

A further challenge is generating database queries that flexibly interface with the logic of neuroimaging and processing pipelines, which is relevant in the context of missing data. `HALFpipe` always tries to execute the best possible processing pipeline based on the data that is available. For example, a field map may have been routinely acquired before each functional scan in a particular dataset. If one of these field maps is missing, `HALFpipe` flexibly assigns another field map, for example one belonging to the preceding functional scan. However, `HALFpipe` will not use a field map from another scan session, as field inhomogeneities are likely to have changed. Finally, `HALFpipe` does not fail if a field map is missing, but simply omits the distortion correction step for that subject. Other examples include the ability of `HALFpipe` to match structural to functional images, and match task events to a functional scan. This strategy is used throughout the construction of processing workflows.

### Metadata

Processing of neuroimaging data requires access to relevant metadata, such as temporal resolution, spatial resolution, and many others. Some elements of metadata, such as echo time (TE), are represented differently depending on scanner manufacturer and DICOM conversion software. The method for reading various types of data has been harmonized in `HALFpipe` using the following three methods.

First, metadata can be stored in BIDS format. This means that a JavaScript Object Notation (JSON) file accompanies each image file, which contains the necessary metadata. BIDS calls this file the *sidecar*, and common tools such as `heudiconv` (Halchenko et al., 2018) or `dcm2niix` (Li et al., 2016) generate these files automatically. If these files are present, `HALFpipe` will detect and use them. Second, instead of sidecar files, some software tools store image metadata in the NIf TI header. The NIf TI format defines fields that can fit metadata, but depending on how the image file was created, these metadata may be missing. Some conversion programs also place the metadata in the description field in free text format. This description can also be parsed and read automatically. Third, information may be incorrectly represented due to user error, incompatible units of measurement, or archaic technical considerations. In such cases, `HALFpipe` provides a mechanism to override the incorrect values. For every metadata field, the user interface will prompt the user to confirm that metadata values have been read or inferred correctly. The user can choose to manually enter the correct values.

### Interfaces

`HALFpipe` consists of different modules that need to pass data between each other, such as file pathnames and the results of quality assessment procedures. Developing an application as large and complex as `HALFpipe` requires establishing predictable interfaces, which prescribe data formats for communication within the application. An advantage of this approach is that knowledgeable users can write their own code to interface with `HALFpipe`.

`HALFpipe` uses the Python module `marshmallow` to implement interfaces, called schemas in the module’s nomenclature. All schemas are defined in the `HALFpipe` code. When the user first starts the application, the user interface is displayed by `HALFpipe`. It asks the user a series of questions about the data set and the analysis plan, and stores the inputs in a configuration file called `spec.json`. The configuration file has a predictable syntax and can be easily scripted or modified, which enables collaborative studies to harmonize analysis plans.

### Workflow engine

To obtain reproducible results, a core requirement for `HALFpipe` was reproducible execution of the processing pipeline. As the ENIGMA consortium requires fMRI analysis of large datasets with several thousand samples, `HALFpipe` was designed to parallelize processing on multiple computers or processor cores. Both of these specifications were achieved by implementation in `Nipype`, *NeuroImaging in Python:Pipelines and Interfaces* (Gorgolewski et al., 2011). `Nipype` is a workflow engine for neuroimaging that constructs an acyclic directed graph, in which nodes represent processing commands that need to be executed (the steps of the pipeline), while the edges represent inputs and outputs being passed between nodes (images or text files). In this formalization of a neuroimaging pipeline as a graph, the fastest order for execution across multiple processor cores can be determined.

The workflow graphs are modular and scalable, which means they can be nested and extended. `HALFpipe` uses the workflows defined by `fMRIPrep` and then connects their outputs to additional workflows. `fMRIPrep` itself is modular and divided into multiple workflows: `sMRIPrep` (Esteban et al., 2021), `SDCFlows`, and `NiWorkflows`. The workflow graph facilitates saving and verifying intermediate results, and supports the user’s ability to stop and later restart processing. `HALFpipe` also uses the graphs to determine which intermediate results files are not needed by subsequent commands by using a tracing garbage collection algorithm (Dijkstra et al., 1978). As such, intermediate files do not accumulate on the storage device. This feature is implemented as a plugin to `Nipype`.

`Nipype` forms the basis of `fMRIPrep` and `C-PAC`, which are widely used in the neuroimaging community. However, it has several limitations that are relevant in the context of `HALFpipe`. `HALFpipe` is able to calculate features and statistical maps with different variations of preprocessing settings. To do this efficiently, intermediate results need to be re-used whenever possible. An improved second version of `Nipype` is currently being developed, called `Pydra` (Jarecka et al., 2020), which will be able to automatically detect repetitive processing commands, and automatically re-use outputs. Presently, until `Pydra` becomes available, `HALFpipe` calculates a four-letter hash code that uniquely identifies each preprocessing step. Before constructing a new preprocessing command, `HALFpipe` checks whether its hash has already been added to the graph. If present, the existing command is re-used.

A key requirement of `HALFpipe` was robust and flexible handling of missing data. For instance a missing functional scan or statistical map does not cause `HALFpipe` to fail. Additionally, `HALFpipe` defines inclusion and exclusion criteria for scans, such as the maximum allowed motion (mean framewise displacement) or a minimum brain coverage when extracting a brain region’s average signal. Finally, depending on the data set, statistical maps may need to be aggregated across runs or sessions within single subjects before a group-level model can be run. This means that the static graph has to be modified dynamically to adapt to the results of processing. `HALFpipe` solves this problem by defining a data structure that not only contains the file names of statistical maps, but also the tags and metadata that can be used to adjust processing on the fly. For example, using this data structure, design matrices can be constructed for group models based on the actual subjects that have statistical maps available.

### Running on a high-performance cluster

`HALFpipe` provides a simple way to run on a high-performance compute cluster. For each subject, preprocessing and feature extraction takes 6 to 10 hours on a single processor core. Most jobs take about 8 GB to 20 GB of RAM, depending on size of the data. Assuming each subject is assigned to a unique job, we recommend requesting 24 GB of RAM for each job. The most memory-intensive steps of processing are spatial registration and resampling, and running `FSL MELODIC` for independent component analysis (ICA) as part of `ICA-AROMA` (Pruim et al., 2015). For large datasets, parallel processing of subjects is highly desirable to reduce the total computation time.

Deploying `Nipype` to perform computations on multiple nodes, such as on a high performance cluster (HPC) is particularly challenging. By default, `Nipype` submits a separate job to the cluster queue for each processing command (graph node) regardless of the amount of time required to execute the command. A watcher process running on the head node collects outputs from completed commands and submits the next processing command. This process can be inefficient on some HPCs because computational resources need to be allocated and deallocated continually. We implemented a more efficient approach for `HALFpipe` that partitions the processing graph into many independent subgraphs, which the user may submit as separate jobs. The smallest granularity available is one subgraph per subject that is invoked with the command line flag `--subject-chunks`. A `Nipype` workflow is created and validated for all subjects before the pipeline starts running. In a cluster setting, the most efficient resource utilization is to submit each subject as a separate job and to run each job on two CPU cores.

### Data denoising

`HALFpipe` and `fMRIPrep` are modular preprocessing pipelines, meaning that they utilize a series of tools from various software libraries. Most have been adopted as standard practice by the community for many years. Thus, the reasons motivating specific algorithmic choices may not be readily apparent to users, but need to be considered when designing tools such as `HALFpipe`. Here we articulate the design considerations in our selection of tools.

`HALFpipe` performs all denoising in a predefined order after resampling the fMRI data to standard space using `fMRIPrep`. `HALFpipe` defines standard space as the *MNI152NLin2009cAsym* template, which is the most current and detailed template available (Horn, 2016a).

`fMRIPrep` not only outputs a preprocessed image in standard space, but also a spreadsheet with nuisance (or confound) time series. These include (derivatives of) motion parameters (squared), aCompCor components (Behzadi et al., 2007), white matter signal, CSF signal, and global signal. A key consideration is that including these time series as nuisance regressors may re-introduce variance that was already removed in previous processing steps (Hallquist et al., 2013). An example of this phenomenon may be regressing out motion parameters after removing low-frequency drift via temporal filtering. In practice, this means setting up a regression model for each voxel, where the temporally filtered time series of a voxel is the dependent variable and the regressors are the motion parameters. The regression model will calculate a regression weight for each motion regressor, so that the total model explains the maximum amount of variance (under assumption of normality). After multiplying the motion parameters with these weights, they are summed to yield one time series containing the motion-related noise. This time series is subtracted from the temporally filtered voxel time series to yield the result of the procedure, the denoised time series (i.e., the regression residuals). However, if the motion parameters happen to contain any low-frequency drift, then their weighted sum likely will as well. It follows that subtracting a time series with temporal drift from the temporally filtered voxel data will introduce temporal drift again, independent of whether a temporal filter was applied before. In `HALFpipe`, any filter or transformation applied to the voxel time series is also applied to the nuisance time series. This way, previously removed variance is not re-introduced accidentally, because it has been removed from both sides of the regression equation. In `HALFpipe`, denoising is implemented as follows:

1. `ICA-AROMA` noise component classification (Pruim et al., 2015) relies on reference masks defined in *MNI152NLin6Asym* space, which is different from the standard space template used by `fMRIPrep`. To solve this issue, `fMRIPrep` will estimate a second normalization to this template, apply it to the fMRI image in native space, and run `ICA-AROMA` on the resulting image (Ciric et al., 2021). This approach effectively doubles the processor time spent on spatial normalization, and may require manually checking both spatial registrations. To avoid this considerable effort, `HALFpipe` resamples the image to *MNI152NLin6Asym* space using an existing transformation/warp field between the two spaces (Horn, 2016b). Specifically, this predefined warp field is concatenated with the subject’s warp to *MNI152NLin2009cAsym* space, with which resampling is performed with `fMRIPrep`’s `bold_std_trans_wf` workflow. Finally, `ICA-AROMA` is run on the resulting fMRI image in *MNI152NLin6Asym* space using `fMRIPrep`’s `ica_aroma_wf` workflow, which also includes spatial smoothing fixed to a 6 mm FWHM smoothing kernel (Pruim et al., 2015). `ICA-AROMA` generates a set of component time series and a binary classification of these components as either signal or noise.
2. `HALFpipe` implements spatial smoothing using `AFNI`’s `3dBlurInMask` (Cox, 1996). Each voxel’s signal is averaged with the signal of its neighboring voxels, weighted by an isotropic gaussian kernel. At the edges of the brain, this kernel may include non-brain voxels, so smoothing is constrained to only happen within the brain mask. This is equivalent to the procedure in the *Minimal Preprocessing Pipelines for the Human Connectome Project* (Glasser et al., 2013).
3. Grand mean scaling sets the image mean, defined as the within-scan mean across all voxels and time points, to a predefined value. The grand mean is closely related to scanner parameters such as amplifier gain but not to neural mechanisms (Gavrilescu et al., 2002). Adjusting the grand mean via scaling makes analysis results more interpretable and comparable across subjects, sessions, and sites. The scaling factor is calculated based on the masked functional image, and applied to both the fMRI data and the nuisance time series extracted by `fMRIPrep`.
4. The previously estimated `ICA-AROMA` noise components are removed from the smoothed and grand-mean-scaled fMRI data. This is done in a non-aggressive way to minimize removing variance that is shared between signal and noise components. ICA-AROMA implements this step using the `FSL` command `fsl_regfilt`, which calculates an ordinary least squares regression for each voxel, where the design matrix includes both the signal and the noise components as regressors. This means that the resulting regression weights reflect the unique variance of the noise components (and not the shared variance with signal components). Then, the noise component regressors are multiplied by their regression weights and added together to yield one time series of all the noise. Subtracting the noise from the voxel time series yields a denoised time series (the regression residuals); this step is done using a re-implementation of `fsl_regfilt` in `Numpy` (Harris et al., 2020). The same procedure is applied to the nuisance time series from step 3.
5. Temporal filtering can be used to remove low-frequency drift via a high-pass filter, high-frequency noise via a low-pass filter, or both at the same time using a band-pass filter. `HALFpipe` implements two approaches to temporal filtering, a frequency-based approach (Jo et al., 2013) and a Gaussian-weighted approach (Marchini & Ripley, 2000). The frequency-based temporal filter is very exact in selecting frequencies to be kept or removed, and is commonly used to calculate fractional Amplitude of Low Frequency Fluctuations (fALFF) and Regional Homogeneity (ReHo). The Gaussian-weighted temporal filter is the default used by `FSL Feat` (Jenkinson et al., 2012) and may have fewer edge effects at the start and end of the time series. However, its spectrum also has a more gradual roll-off, meaning that it will be less aggressive in removing frequencies close to the chosen cutoff value. Temporal filtering is applied to both the fMRI data and nuisance time series from step 4.
6. Nuisance time series from step 5 are removed using the regression residualization procedure described above from both the fMRI data and the nuisance time series.

`HALFpipe` suggests default settings for each of these steps, which are outlined in Table 2. Note that some are selected based on best-practices in the field (i.e., band-pass temporal filter for ALFF and ReHo), whereas most default settings can be adjusted by the user.

### Quality assessment

Assessing the quality of data and preprocessing is a laborious undertaking and often done manually. Efforts to automate this process, either through predefined thresholds of image quality features (Alfaro-Almagro et al., 2018) or machine learning (Esteban et al., 2017) are not yet ready to replace the eyes of a trained researcher checking the data. However, various approaches make this process easier. First, rather than viewing three-dimensional neuroimaging files directly, generating and viewing reports containing two-dimensional images offers a significant time savings. Second, tools such as `slicesdir` (in `FSL`), `fMRIPrep`, and `MRIQC` generate HTML files that contain multiple report images and can be explored in a web browser. `MRIQC` also provides an interactive widget to rate the quality of each image (Esteban, Blair, et al., 2019).

In `HALFpipe`, we use a fixed set of processing steps for quality assessment. While `slicesdir` allows the researcher to easily compare the same image type across different subjects, it cannot be used to generate reports for all types of images. By contrast, `fMRIPrep` / `MRIQC` HTML files have a broad range of information and quality report images included, but one HTML file is always specific to one subject. As such, examining multiple processing steps in many subjects can be cumbersome. To overcome these issues, `HALFpipe` provides an interactive web app that is contained in a single HTML file. The app dynamically loads reports with images, and can handle datasets up to thousands of images without a performance penalty. The images can be sorted both by subject, as is done by `fMRIPrep` / `MRIQC`, or by image type, as is done in `slicesdir`. Each image can be rated as either good, uncertain, or bad. Predefined logic automatically converts these ratings into inclusion/exclusion decisions for `HALFpipe`’s group statistics. In addition, tagging images as uncertain enables users to efficiently retrieve and discuss these with a colleague or collaborator, after which a definitive decision on image quality can be made.

Images can be zoomed by clicking them. For faster operation by advanced users, rating and navigation are accessible not just via user interface buttons, but also via keyboard shortcuts based on the WASD keys. Pressing the A goes back one image and D goes ahead, whereas W, S and X rate an image as good, uncertain or bad, respectively. The web app offers an overview chart that indicates subjects preprocessed successfully and subjects with errors, a chart with quality ratings, and box plots reflecting the sample distributions for motion, noise components, and temporal signal-to-noise ratio (tSNR). All three are implemented so that users can hover over chart elements with their cursor to view meta-information, such as the subject identifier, and click to navigate to the associated report images. The HTML file is built as a frameworkless web app using TypeScript. Source code is available at https://github.com/HALFpipe/QualityCheck.

`HALFpipe` shows two report images for each subject on structural/anatomical processing and four additional images for each type of functional scan. Detailed explanations may be found in the quality assessment manual at https://github.com/HALFpipe/HALFpipe#quality-checks.

1. T1w skull stripping shows the bias-field corrected anatomical image overlaid with a red line that outlines the brain mask. The user must check that no brain regions are missing from the mask, and that portions of the skull or head are not included in the mask.
2. T1w spatial normalization shows the anatomical image resampled to standard space overlaid with a brain atlas in standard space. The user needs to check whether the regions of the atlas closely match the resampled image.
3. Echo planar imaging (EPI) tSNR shows the temporal signal-to-noise ratio of the functional image after preprocessing using `fMRIPrep`. The user must check that signal recovery is distributed uniformly throughout the brain, and exclude scans with asymmetry, distortions, localized signal drop-out, or striping artifacts.
4. EPI Confounds shows the carpet plot (Aquino et al., 2020; Power, 2017) generated by `fMRIPrep`. A carpet plot is a two-dimensional plot of time series within a scan, with time on the *x*-axis and voxels on the *y*-axis. Voxels are grouped into cortical gray matter (blue), subcortical gray matter (orange), cerebellum (green), and white matter and cerebrospinal fluid (red). Above the carpet plot are time courses (x-axis) of the magnitude (y-axis) of framewise displacement (FD), global signal (GS), global signal in CSF (GSCSF), global signal in white matter (GSWM), and DVARS, which is the temporal change in root-mean-square intensity (D = temporal derivative of time courses, and VARS = root-mean-square variance over voxels). The user must look for changes in heatmap/intensity in relation to motion and signal changes above. Abrupt changes in the carpet plot may correspond to motion spikes, whereas extended signal changes may indicate acquisition artifacts caused by defective scanner hardware.
5. EPI ICA-based artifact removal shows the time course of the mean signal extracted from each ICA-component and its classification as either signal (green) or noise (red). This figure is generated by `fMRIPrep`. For each component, there is a spatial map (left), the time series (top right) and the power spectrum (bottom right). The user must check that components classified as noise do not contain brain networks or temporal patterns that are known to be signal.
6. EPI spatial normalization shows the functional image after preprocessing using `fMRIPrep` overlaid with a brain atlas in standard space. As before, the user must check whether the regions of the atlas closely match the resampled image.

### Statistics

`HALFpipe` uses `FSL` FMRIB Local Analysis of Mixed Effects (FLAME) (Woolrich et al., 2004) for group statistics, because it considers the within-subject variance of lower level estimates in its mixed effects models. In addition, its estimates are conservative, which means they offer robust control of the false positive rate (Eklund et al., 2016).

A common issue in fMRI studies is that the spatial extent of brain coverage may differ between subjects. A common choice is to restrict higher-level statistics to only those voxels that were acquired in every subject. However, with a large variation in brain coverage, which is to be expected when pooling multi-cohort data, sizable portions of the brain may ultimately be excluded from analysis. To circumvent this issue, `HALFpipe` uses a re-implementation of `FSL`’s `flameo` in `Numpy` (Harris et al., 2020). In this implementation, a unique design matrix is re-generated for every voxel so that only subjects who have a measurable value for a given voxel are included. Then the model is estimated using the FLAME algorithm. This list-wise deletion approach depends on the assumption that voxels are missing completely at random (MCAR), meaning that the regressors (and thus statistical values) are independent of scanner coverage.

For group models, users can specify flexible factorial models that include covariates and group comparisons. By default, missing values for these variables are handled by list-wise deletion as well, but the user may alternatively choose to replace missing values by zero in the demeaned design matrix. The latter approach is equivalent to imputation by the sample mean. Design matrices for the flexible factorial models are generated using the Python module `Patsy` (Smith et al., 2018). Contrasts between groups are specified using the `lsmeans` procedure (Lenth, 2016).

## Procedure

`HALFpipe` starts up as a terminal-based user interface that prompts the user with a series of questions about the dataset being analyzed and the desired analysis plan. The main stages of `HALFpipe` analysis, which are detailed below, include loading data, preprocessing with `fMRIPrep`, quality assessment, feature extraction, and group-level statistics. Users have the flexibility to specify the settings for each processing stage at one time or separately at each stage. If `HALFpipe` is stopped and resumed at an intermediate stage, `HALFpipe` will detect which stages have been completed and ask the user to indicate further analyses that are desired. For instance, the user can request preprocessing and feature extraction, but not group-level statistics, and later resume processing specifying group-level statistics only.

### Loading data

A major advantage of `HALFpipe` is that it accepts input data organized in various formats without the need for file naming conventions or a specific directory structure. Using the terminal interface, the user is asked to provide the location of the T1-weighted and fMRI BOLD image files, which are required for preprocessing, as well as field maps and task event files if available or applicable. However, `HALFpipe` requires additional information linking the image files to run in an automated fashion, such as information specifying which set of images belong to the same subject.

Through the use of path templates, `HALFpipe` can handle a wide range of folder structures and data layouts. The syntax for path templates is adapted from `C-PAC`’s data configuration (Giavasis et al., 2020). Instead of manually adding each input file for each subject separately, as is done in the `SPM` or `FSL` user interfaces, the template describes the pattern used for naming files. That pattern can match many file names, thereby reducing the amount of manual work for the user. For example, when placing the tag `{subject}` in the file path `{subject}_t1.nii.gz`, all files of which the name ends in `_t1` and have the extension `.nii.gz` will be selected. The part of the filename that comes before `_t1` is now interpreted by the parsing algorithm as the subject identifier. When multiple files from different modalities have the same subject identifier, or session number, etc., they will be matched automatically by these tags. Automated processing workflows can then be constructed around the resulting data structure.

In contrast to `C-PAC`’s data configuration syntax, `HALFpipe` path templates use BIDS tags (Gorgolewski et al., 2016). `HALFpipe` path templates can be further specified by adding a colon and a regular expression after the tag name (as in standard Python regular expression syntax). For example, `{subject:[0-9]}` will only match subject identifiers that contain just digits. This can be useful for more complex data layouts, such as when multiple datasets are placed in the same directory, and only a single subset is to be used. For more examples, see Table 3.

**Table 3:** Examples of path template syntax

|  | Example 1 | Example 2 |
| --- | --- | --- |
| Path template | data/{subject}/bold_rest.nii.gz | data/{subject}/bold_{task}.nii.gz |
| Matches these files | data/subject01/bold_rest.nii.gz<br>data/subject02/bold_rest.nii.gz<br>data/subject03/bold_rest.nii.gz<br>data/phantom/bold_rest.nii.gz | data/subject01/bold_rest.nii.gz<br>data/subject02/bold_rest.nii.gz<br>data/subject03/bold_rest.nii.gz<br>data/subject01/bold_task.nii.gz |
| Does not match | data/subject01/bold_task.nii.gz<br>data/subfolder/subject01/bold_rest.nii.gz | data/phantom/bold_rest.nii.gz<br>data/subfolder/subject01/bold_rest.nii.gz |

Note: The grey highlight shows the part of the file name that is responsible for not matching.

In the `HALFpipe` user interface, the user receives feedback on how many and which files are matched, so that the path templates can be entered interactively. Importantly, after finishing the configuration process via the user interface, all files are internally converted into the standardized BIDS structure, which is a prerequisite for running `fMRIPrep`. However, no copies of files are made, the conversion is based entirely on symbolic links (aliases) to the original files. If the data are already in BIDS format, `HALFpipe` will still carry out this conversion for consistency. The resulting dataset in BIDS format is then stored in the working directory in a subfolder called `rawdata`.

### Preprocessing

Preprocessing is implemented in `HALFpipe` using `fMRIPrep`, which performs a consensus of preprocessing steps required for any fMRI study (Esteban, Markiewicz, Blair, et al., 2019). The consensus steps include skull stripping, tissue segmentation, and spatial normalization of structural images. Consensus steps for functional images include motion correction, slice time correction, susceptibility distortion correction, and spatial normalization. Besides slice timing correction, all other steps entail spatial transformations. `fMRIPrep` calculates the parameters for each of these transformations, which are combined and finally applied in one step. The resulting image is outputted, among other derived data and a report on preprocessing. Importantly, `HALFpipe` runs `fMRIPrep` with small modifications. We disabled experimental susceptibility distortion correction in the absence of field maps, because it is not yet validated. We also do not output preprocessed and normalized functional images by default, because they use a lot of disk space. However, users can manually choose to output preprocessed images with their choice of preprocessing settings in the user interface.

For fMRI data, `HALFpipe` can perform denoising via `ICA-AROMA` (Pruim et al.,2015). Additional widely used preprocessing steps, such as spatial smoothing, grand mean scaling, temporal filtering (Gaussian-or frequency-based), and confound regression can be selected by the user, the latter using confounds selected from the large set generated by `fMRIPrep`, including the original motion parameters, derivatives of motion parameters, motion parameters squared, top five aCompCor components (Behzadi et al., 2007), white matter signal, CSF signal, and global signal. Importantly, in `fMRIPrep` all confound signals are extracted from the data before `ICA-AROMA` is run. If applicable, `HALFpipe` will apply denoising to these confound signals as well, to match the preprocessed and denoised data, so as to not accidentally re-introduce noise variance (Hallquist et al., 2013; Lindquist et al., 2019). This was illustrated with the example on motion parameter regression in the section Data denoising.

Various confound and denoising settings may be used for each fMRI feature (see section Feature extraction), and for generating preprocessed images that can, for example, be used to extract features with software other than `HALFpipe`.

### Quality assessment

Quality assessment can be performed in an interactive, browser-based user interface (see ***Figure 2***). `HALFpipe` provides a detailed user manual for quality assessment that is linked on the web page. The web app shows report images of several preprocessing steps such as T1 skull stripping and normalization, BOLD tSNR, motion confounds, ICA-based artifact removal, and spatial normalization (see the methods section on Quality assessment). These images can be visually inspected and rated by the viewer as either good, uncertain, or bad.

**Figure 1.**
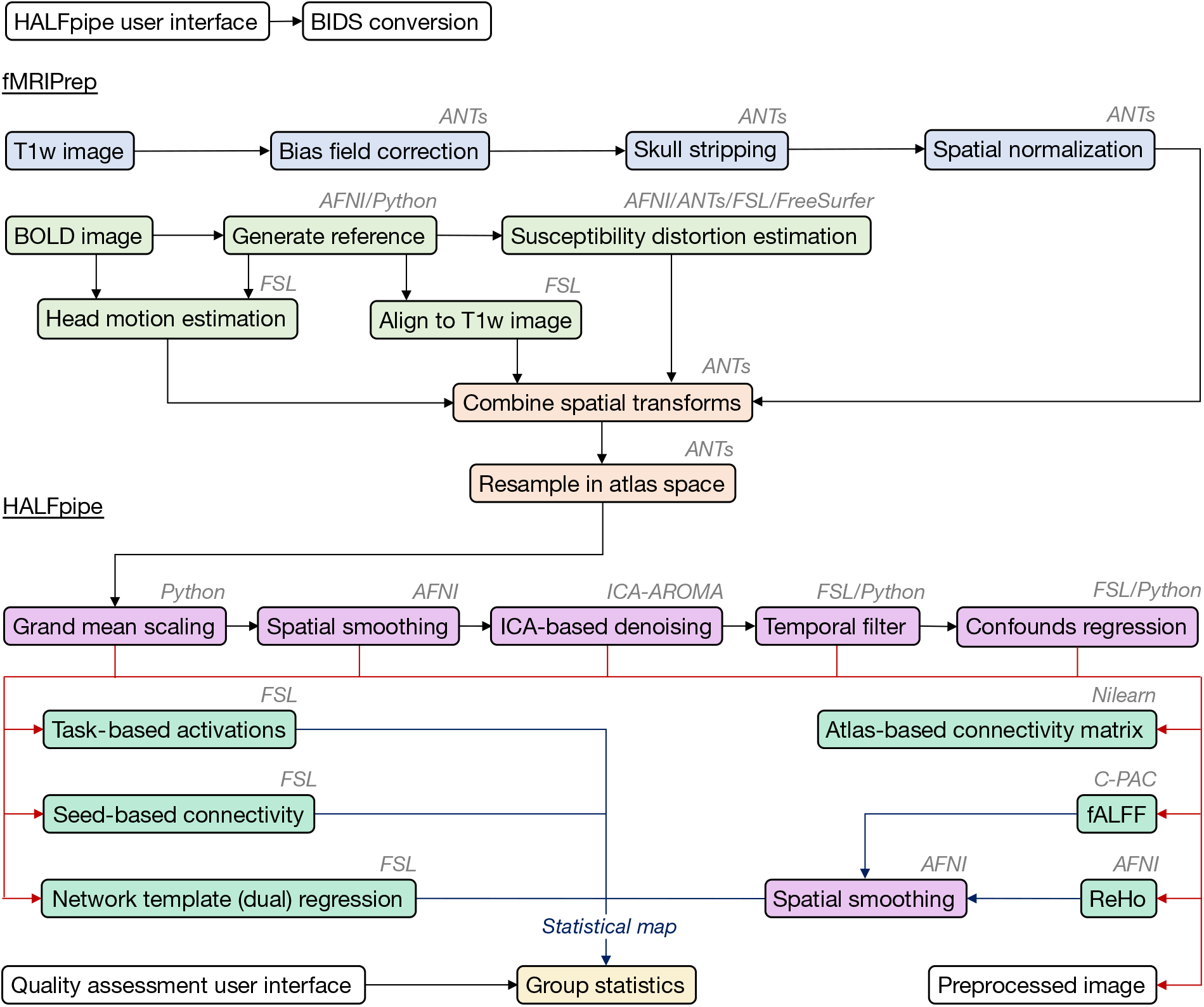
HALFpipe workflow. After minimal preprocessing with fMRIPrep (Esteban, Markiewicz, Blair, et al., 2019), additional preprocessing steps can be selected (purple). Using the preprocessed data, statistical maps can be calculated during feature extraction (turquoise). Note that not all preprocessing steps are available for each feature, as is outlined in Table 2. The diagram omits this information to increase visual clarity.

**Figure 2.**
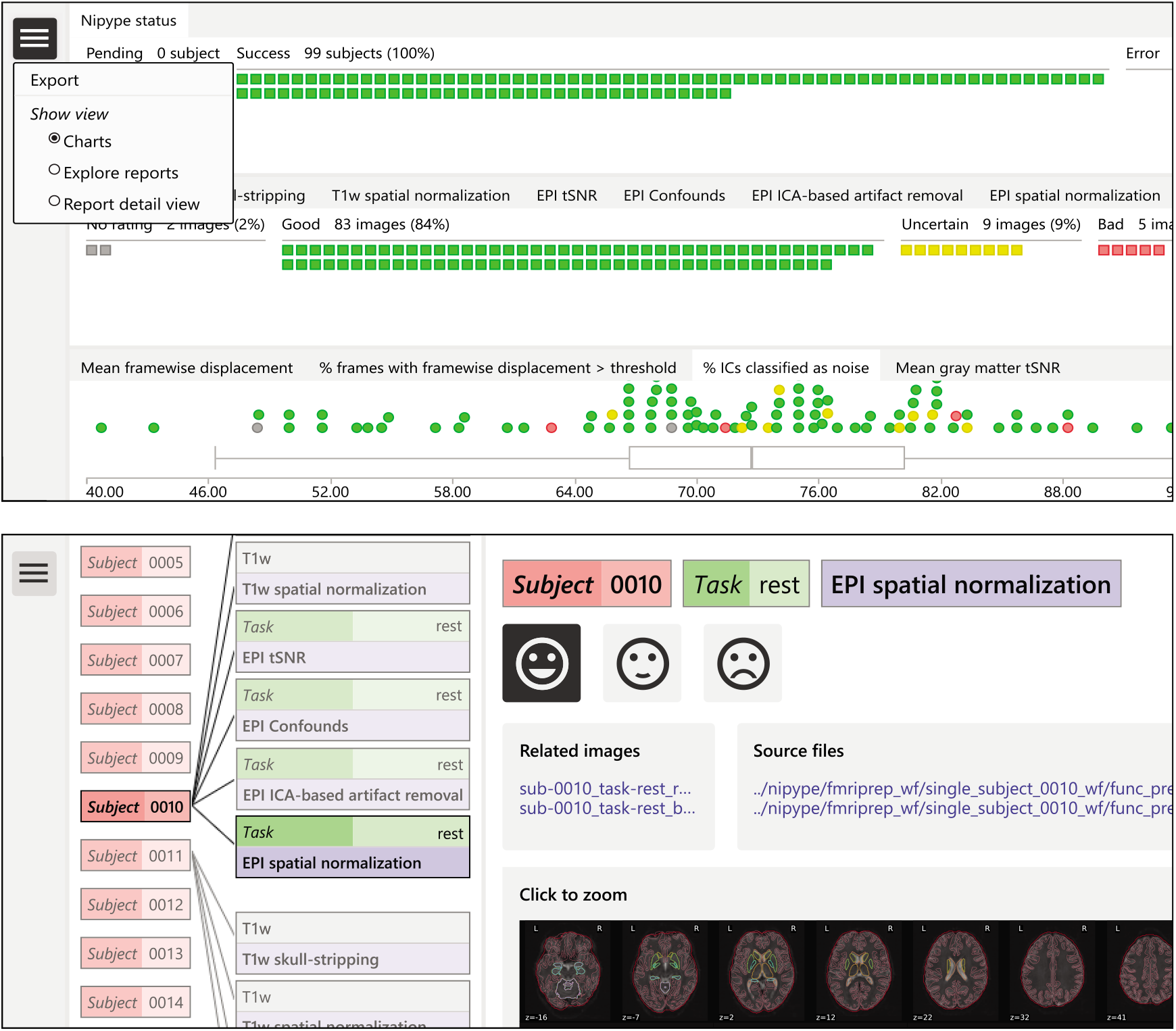
Quality assessment user interface. The top panel shows the charts view, containing one chart for processing status, one for quality ratings and one for image quality metrics. In the top left corner, the navigation menu is open, which shows the option to export ratings for use in group statistics. The bottom panel contains a screenshot of the explorer view that allows the user to navigate across subjects and image types. The explorer view shows the currently selected report image on the right, along with its rating, related images, and the source files that were used to construct it.

Ratings will be saved in the local browser storage. Once completed, they can be downloaded in JSON format to be read by `HALFpipe`. If placed in the working directory, ratings will be automatically detected by `HALFpipe` and used to exclude subjects for group-level statistics. Additionally, `HALFpipe` will automatically detect all other JSON files whose names start with `exclude`, to accommodate quality assessment by multiple researchers. In the case of conflicts between ratings, the lower rating will be used.

`HALFpipe` will include as much data as possible while excluding all scans rated as “bad”. Ratings of “good” and “uncertain” will be included for group analysis. A “bad” rating for any report image related to structural/anatomical processing will exclude the entire subject. A “bad” rating for any report image related to functional image processing will only exclude the specific functional scan. This means that if a subject has one “bad” scan, its other scans may still be included for group statistics.

In addition, the mean framewise displacement, percentage of frames with a framewise displacement above a specified threshold, percentage of the independent components that were classified as noise, and mean gray matter tSNR from all subjects is displayed in box plots. Next to the report images, links to the source images are shown so that these can be inspected in more detail by opening them in a preferred image viewer (e.g., `fsleyes`).

### Feature extraction

Following preprocessing, `HALFpipe` can extract several *features* that are commonly used in resting-state and task-based analysis. These include various ways of examining functional connectivity between brain regions (seed-based connectivity, network-template (or dual) regression, atlas-based connectivity matrices), as well as measures of local activity (ReHo, fALFF). `HALFpipe` allows the user to choose several region-of-interest masks (seeds), template networks, and atlases, for which a threshold indicates the minimum overlap the user requires between seeds, template networks, or atlas regions and the subjects’ fMRI data. For each feature, the user can change the default settings for spatial smoothing and temporal filtering, and choose the confounds to be removed. The user is offered the option to extract the same feature multiple times, each time varying the preprocessing, confound, and denoising settings to explore the impact of analytical decisions in a *multiverse analysis*. Of note, for selected features some options are not available. For example, spatial smoothing is disabled for atlas-based connectivity matrices (Alakörkkö et al., 2017), or performed *after* ReHo and fALFF have been calculated (see Table 2).

A brief description of the features is provided in Box 1.

#### Box 1: Overview of HALFpipe features

##### Task-based activations

A first-level general linear model (GLM) is run for event-related or block designs. GLM regressors describing the stimulus presentations for each of the task conditions are convolved with a double Gamma HRF and the overall model is fit for each voxel in the brain using `FSL FILM` (Woolrich et al., 2001). Contrasts of interest are tested, which results in a whole-brain task activation map for comparisons between task conditions.

##### Seed-based connectivity

Average BOLD time series are extracted from a region of interest (seed), which is defined by a binary mask image. This time series is used as a regressor in a first-level GLM, where the model is fit for each voxel in the brain using `fsl_glm`. This results in a whole-brain functional connectivity map that represents the connectivity strength between the ROI and each voxel in the brain.

##### Network-template (or dual) regression

Subject-specific representations of connectivity networks (e.g., default mode, salience, task-positive networks) are generated using dual regression (Beckmann et al., 2009) with `fsl_glm`. In a first regression model, the set of network template maps is regressed against the individual fMRI data, which generates time series for each of the template networks. Next, a second regression model is run, regressing the network time series against the individual fMRI data. This generates subject-specific spatial representations of each of the template networks, which can be considered to represent the voxelwise connectivity strength within each of the networks.

##### Atlas-based connectivity matrix

Average time series are extracted from each region of a brain atlas of choice using custom code inspired by `Pypes` (Savio et al.,2017)and `Nilearn` (Abraham et al., 2014). From these, a pairwise connectivity matrix between atlas regions is calculated using Pearson product-moment correlations using `Pandas` (McKinney, 2010), which represent the pairwise functional connectivity between all pairs of regions included in the atlas.

##### Regional homogeneity (ReHo)

Local similarity (or synchronization) between the time series of a given voxel and its nearest neighboring voxels is calculated using Kendall’s coefficient of concordance (Zang et al., 2004) using `FATCAT`’s `3dReHo` which is distributed with `AFNI` (Taylor & Saad, 2013).

##### Fractional amplitude of low frequency fluctuations (fALFF)

Variance in amplitude of low frequencies in the BOLD signal is calculated, dividing the power in the low frequency range (0.01–0.1 Hz) by the power in the entire frequency range (Zou et al., 2008) with a customized version of the `C-PAC` implementation of fALFF.

### Group-level statistics

Group-level statistics on individual features can be performed with `FSL`’s FLAME algorithm. Subjects who had poor quality data in the interactive quality assessment are excluded. In addition, subjects can be excluded based on movement by selecting the maximum allowed mean framewise displacement (FD) and percentage of outlier frames (i.e., frames with motion higher than the specified FD threshold).

For group-level statistics, users can choose to calculate the intercept only (i.e., mean across all subjects) or run flexible factorial models. For the latter, `HALFpipe` prompts the user to specify the path to a covariates file (multiple file formats are supported) containing subject identifiers, group membership, and other variables, and to specify whether these are continuous or categorical. Missing values in the covariates file can be handled with either listwise deletion or mean substitution. The user can specify main effects and interactions between variables, while within-group regressions against a continuous variable (e.g., symptom severity) is also possible.

## Discussion

Large samples are essential for recent neuroimaging applications, such as imaging-genetics association studies, training of complex machine learning models, and even unsupervised learning. This demand has stimulated efforts to pool data from multiple observational studies, which typically incur greater bias than studies designed *a priori* to address a specific scientific question. Within ENIGMA, we developed `HALFpipe` to support harmonization of task-based and resting-state fMRI data analysis and quality assessment across multiple labs and cohorts. `HALFpipe` bundles all software tools, library functions, and other dependencies by containerizing the requisite components in a Singularity (Kurtzer et al., 2017) and Docker (Docker Inc.) release. Containerization ensures that all software dependencies and the runtime environment are provided. Therefore, containerized software such as `HALFpipe` can run reliably regardless of the computing environment where it is installed, be it a laptop, computational cluster, or cloud computing service (Grüning et al., 2018).

The design, implementation, and testing of the `HALFpipe` workflow resulted in its 1.0 version release in early 2021. Several thousand resting-state fMRI datasets from 29 ENIGMA PTSD consortium sites have already been analyzed as part of the first published report to employ `HALFpipe` (Weis, 2020), while analyses of other large multi-site datasets are currently underway in several ENIGMA working groups, including the ENIGMA task-based fMRI working group (Veer et al. 2019). Running `HALFpipe` requires approximately 8 to 20 GB of RAM per computer or cluster node and 6 to 10 hours to complete on a single processor core. The exact resource usage depends on voxel resolution and the number of volumes in the fMRI data. The number of features the user chooses has a negligible impact on processing time.

The `HALFpipe` user experience includes an interactive user interface to facilitate rapid analysis prototyping while preserving the ability to script automated analyses of large datasets via configuration files in JSON format with detailed prescriptions of the dataset, analyses steps, and input parameters. Importantly, `HALFpipe` accommodates concurrent harmonized processing of task-based and resting-state fMRI data, which facilitates cross-modal comparisons between the two fMRI modalities.

Our implementation of `HALFpipe` enables users to tackle consortium analyses of multicohort fMRI data with highly uniform application of methods. Specifically, we have established a standardized process and analysis methodology that involves a pre-specified: (1) ensemble of software tools, (2) software version for each tool, (3) set of user-defined parameters, (4) analytic steps, (5) sequence of analytic steps, (6) quality assessment process, and (7) criteria for excluding substandard data. Thus, `HALFpipe` promotes the seamless implementation of a standardized process (preprocessing and feature extraction) at each site and/or cohort prior to initiating group level statistics. Such capabilities hold the promise of significantly advancing basic neuroscience, and particularly clinical neuroscience, by supporting the execution of multi-site multi-cohort studies of several hundred or several thousand samples — ultimately supporting harmonized cross-disorder comparisons. While not part of the `HALFpipe` workflow, cross-site/platform harmonization techniques for neuroimaging have recently experienced a dramatic increase (Fortin et al., 2018; Pezoulas et al., 2020; Wachinger et al., 2021). Much of this methodological innovation has arrived on the heels of earlier developments in crossplatform harmonization of genetic data (Borisov et al., 2019; Haghverdi et al., 2018; Johnson et al., 2007; Pontikos et al., 2017). These advances in harmonization of neuroimaging data are expected to manifest synergy with standardized workflows such as `HALFpipe`, as both elements are essential to large-scale imaging consortium efforts (Thompson, Jahanshad, et al., 2020).

The implementation of quality metrics for fMRI data has been an incremental process that has moved steadily towards establishing empirically-informed best practices. Historically, quality criteria have been applied unevenly across research labs. Recent years have witnessed a heightened awareness about the essential role of applying systematic and principled quality metrics to minimize confounds, for example motion artifacts (Murphy et al., 2013; Power et al., 2012; Power et al., 2014), and widespread fMRI signal deflections (Aquino et al., 2020). Automated quality control methods are being developed and adopted with increasing interest, such as the MRI Quality Control software `MRIQC` (Esteban et al., 2017). `HALFpipe` has adopted parts of the functionality of `MRIQC` with an enhanced user experience that generates quality reports via a web-browser-based interface to facilitate rapid viewing, screening, and selection of individual subject data for inclusion or exclusion. The application of uniform quality assessment procedures is particularly important when mega-analyzing and even meta-analyzing multi-site/scanner data, as is done in ENIGMA. That is, study variables that segregate by site are more likely to lead to confounds without the uniform implementation of quality assessment across sites (e.g. Wachinger et al., 2021). With its harmonized quality procedures, `HALFpipe` aims to minimize such effects.

### Limitations

Computing platforms that are likely to differ between sites are known to introduce subtle differences in output attributable to operating systems and hardware (Gronenschild et al., 2012). Collecting raw multi-site data at one central site prior to `HALFpipe` processing ensures that the same computing platform can be used to process all data. While optimal, this is often not practical due to restrictions on data sharing, even when the data is completely de-identified (i.e., when linking data to protected health or other sensitive information is no longer possible).

`HALFpipe` offers harmonization through uniform processing of fMRI data, but other sources of non-uniformity are beyond its scope. Recent advances in cross-site/platform harmonization may additionally correct for differences in site, scanner hardware, or computation on different processors (Fortin et al., 2018; Pezoulas et al., 2020; Wachinger et al., 2021). Such methods could be applied to extracted `HALFpipe` features, either centralized or through distributed computation using tools such as `COINSTAC` (Plis et al., 2016), to yield results that are potentially more generalizable.

### Conclusion

`HALFpipe` provides a standardized workflow that encompases the essential elements of task-based and resting-state fMRI analyses, builds on the progress and contributions of `fMRIPrep` developers, and extends capabilities beyond preprocessing steps with a diverse set of postprocessing functions. `HALFpipe` represents a major step toward addressing the reproducibility crisis in functional neuroimaging by offering a workflow that maintains details of user options, steps performed in analyses, metadata associated with analyses, code transparency, containerized installation, and the ability to recreate the runtime environment, while implementing empirically-supported best-practices adopted by the functional neuroimaging community.

## Acknowledgements

This work has been supported by a Grant from the German Research Foundation to S.E. and H.W. (DFG ER 724/4-1, WA 1539/11-1). P.M.T. received research grant funding from Biogen, Inc., for research unrelated to this manuscript. L.S. received funding from the National Institutes of Health (NIH RO1 MH117601 and NHMRC Career Development Fellowship 1140764). R.A.M received funding from National Institutes of Health (NIH R01 MH111671). I.M.V. and H.W. received funding from the European Union’s Horizon 2020 research and innovation programme under under Grant Agreement number 777084 (DynaMORE project).

